# Visual Sequence Encoding is Enhanced by Predictable Music Pairing via Modulating Medial Temporal Lobe and Its Connectivity with Frontostriatal Loops

**DOI:** 10.1101/2023.08.01.551506

**Authors:** Yiren Ren, Thackery Brown

## Abstract

Listening to music during cognitive activities, such as reading and studying, is very common in human daily life. Therefore, it is important to understand how music interacts with concurrent cognitive functions, particularly memory. Current literature has presented mixed results for whether music can benefit learning in other modalities. Evidence is needed for what neural mechanisms music can tap into to enhance concurrent memory processing. This fMRI study aimed to begin filling this gap by investigating how music of varying predictability levels influences parallel visual sequence encoding performance. Behavioral results suggest that overall, predictable music enhances visual sequential encoding, and this effect increases with the structural regularity and familiarity of music. fMRI results indicate that during visual sequence encoding, music activates traditional music-processing and motor-related areas, but decreases parahippocampal and striatal engagement. This deactivation may indicate a more efficient encoding of visual information when music is present. By comparing music conditions of different structural predictability and familiarity, we probed how this occurs. We demonstrate improved encoding with increased syntactical regularity, which was associated with decreased activity in default mode network and increased activity in inferior temporal gyrus. Furthermore, the temporal schema provided by music familiarity may influence encoding through altered functional connectivity between the prefrontal cortex, medial temporal lobe and striatum. Overall, we propose that pairing music with learning might facilitate memory by reducing neural demands for visual encoding and simultaneously strengthening the connectivity between the medial temporal lobe and frontostriatal loops important for sequencing information.

**Significance Statement:** There is considerable interest in what mechanisms can be tapped to improve human memory. Music provides a potential modulator, but few studies have investigated music effects on encoding episodic memory. This study used a novel design to examine how music can influence concurrent visual item sequence encoding. We provided neural data to better understand mechanisms behind potential benefits of music for learning. Our results demonstrated predictable music may help guide parallel learning of sequences in another modality. We found that music might facilitate processing in neural systems associated with visual declarative long-term and working memory, and familiar music might modulate reward circuits and provide a temporal schema which facilitates better encoding of the temporal structure of new non-music information.

## Introduction

The ability to remember the temporal order and sequential relationships between items is essential for various human cognitive functions, such as language, motor skills, episodic memory, and navigation. Given its importance, whether and how sequence learning can be facilitated has become an area of considerable interest. Given the pervasive presence of music in daily human life, and its uniquely predictable temporal sequential structure—which humans adeptly comprehend like a language—we sought to investigate if specific types of music can enhance parallel sequence encoding in other modalities.

Existing literature has reported divergent findings, suggesting that the effects of music on cognition may depend on factors such as music duration, genre, and design(Schwartz et al., 2017). Consequently, researchers have become increasingly interested in the question of ‘which types of music yield greater benefits?’. Some evidence suggests music that is predictable (either familiar or syntactically/structurally regular) can better assist cognitive process (Ford et al., 2016; Koelsch et al., 2005). However, few studies have tested whether predictable music can aid parallel memory of information that is non-linguistic or motoric in nature (e.g., could music facilitate remembering a series of events?). To address this gap, this study developed a novel visual sequence learning paradigm paired with concurrent music which was designed with different forms of predictability.

First, we used fMRI to investigate how concurrent music affected neural recruitment during visual sequence encoding compared to a control condition – an isochronous auditory stream. Specifically, we focused on the recruitment of the medial temporal lobe (MTL), given its crucial role in sequential memory and temporal relationship learning(Schendan et al., 2003) and the striatum. The striatum, in particular, could putatively play a dual role: 1) frontostriatal loops collaborate with MTL to support declarative visual sequence memory (e.g., (Brown et al., 2012) and 2) contribute to music rhythmic processing (Grahn & Rowe, 2013). Given its position at the intersection between MTL and prefrontal cortex(Haber & Knutson, 2010), and between music processing, prediction(Cheung et al., 2019), and sequence processing(Yin, 2010), we thus hypothesized that music modulates visual sequential learning in part via engagement of the striatum and its functional connectivity with other areas including the MTL.

Subsequently, our study aimed to test the hypothesis that highly predictable music provides greater memory benefits, and to examine why. One potential mechanism is attention. Music’s rhythmic and predictable nature can guide attention (Johndro et al., 2019). Notably, predictable music can induce neural entrainment which putatively enhances information transfer between brain regions(Vuilleumier & Trost, 2015), and promotes oscillatory synchrony across prefrontal and temporal lobes during regularly-structured music listening and memory encoding(Thaut et al., 2005, 2014). This phase synchronization has been associated with working memory and long term memory encoding performance(Fell & Axmacher, 2011; Schack et al., 2005). Therefore, we hypothesized that encoding paired with “regular” music (as operationalized) would improve sequence encoding, possibly through more efficient processing in working memory/attention related areas including the prefrontal cortex (PFC).

Furthermore, we explored the potential benefits of music predictability on memory through the lens of “schema” effects on learning. Previous studies suggest that novel information associated with prior knowledge can be encoded more effectively, putatively via interactions between PFC and MTL that allow efficient mapping of relationships between new and old information in memory(van Kesteren et al., 2010, 2012). Music is highly schematic and can provide a rhythmic ‘scaffold’ for new sequence learning(Conway et al., 2009; Leman, 2012); through our music manipulations we examined whether a music schema can influence paired learning in another modality. We designed our task to synchronize the music with the visual stimuli, hypothesizing that more-predictable experimental music conditions could provide a known temporal structure that hippocampus-PFC interactions could leverage to assist encoding temporal relationship between visual items. Critically, the results from our design offer evidence that music’s predictable structure may benefit new visual sequence learning by modulating the dynamic interactions between hippocampus and frontrostriatal systems.

## Method

### Participants

Fifty-four healthy participants were recruited from the Georgia Institute of Technology’s volunteer pool. The sample size was determined *a priori* based on a separate behavioral study (in prep.) using the same experiment design. Two participants failed to attend the second day of the task. One participant withdrew due to reported discomfort during the task. Therefore, the final analysis included data from 51 participants (25 females and 26 males) with an average age is 19.81 years old and a standard deviation of 1.6. Out of the 51 participants, 35 underwent fMRI scanning, while the rest 16 participants completed the task with behavioral measures only. All subjects were pre-screened to exclude any hearing disorders or music recognition disability (such as amusia). Participants who volunteered for the fMRI scan were subject to further screening to exclude any magnetic resonance imaging contradictions (e.g., metallic implants) or major neurological disorders. All participants provided informed consent to the procedures approved by the Institutional Review Board of Georgia Institute of Technology.

### Experimental Design

We developed a two-day sequential memory paradigm (See Figure 1.a). For this, we created novel music compositions and abstract visual stimuli so that familiarity and predictability for both types of information could be controlled experimentally. All the music stimuli (8 seconds each) were composed in the style of Western classical music by trained musicians. We developed two types of music predictability based on the music literature that suggests there are two independent mechanisms for music expectation (veridical expectation and schematic expectation) that can interact and compete with each other(Tillmann & Bigand, 2010).

**Figure 1.**
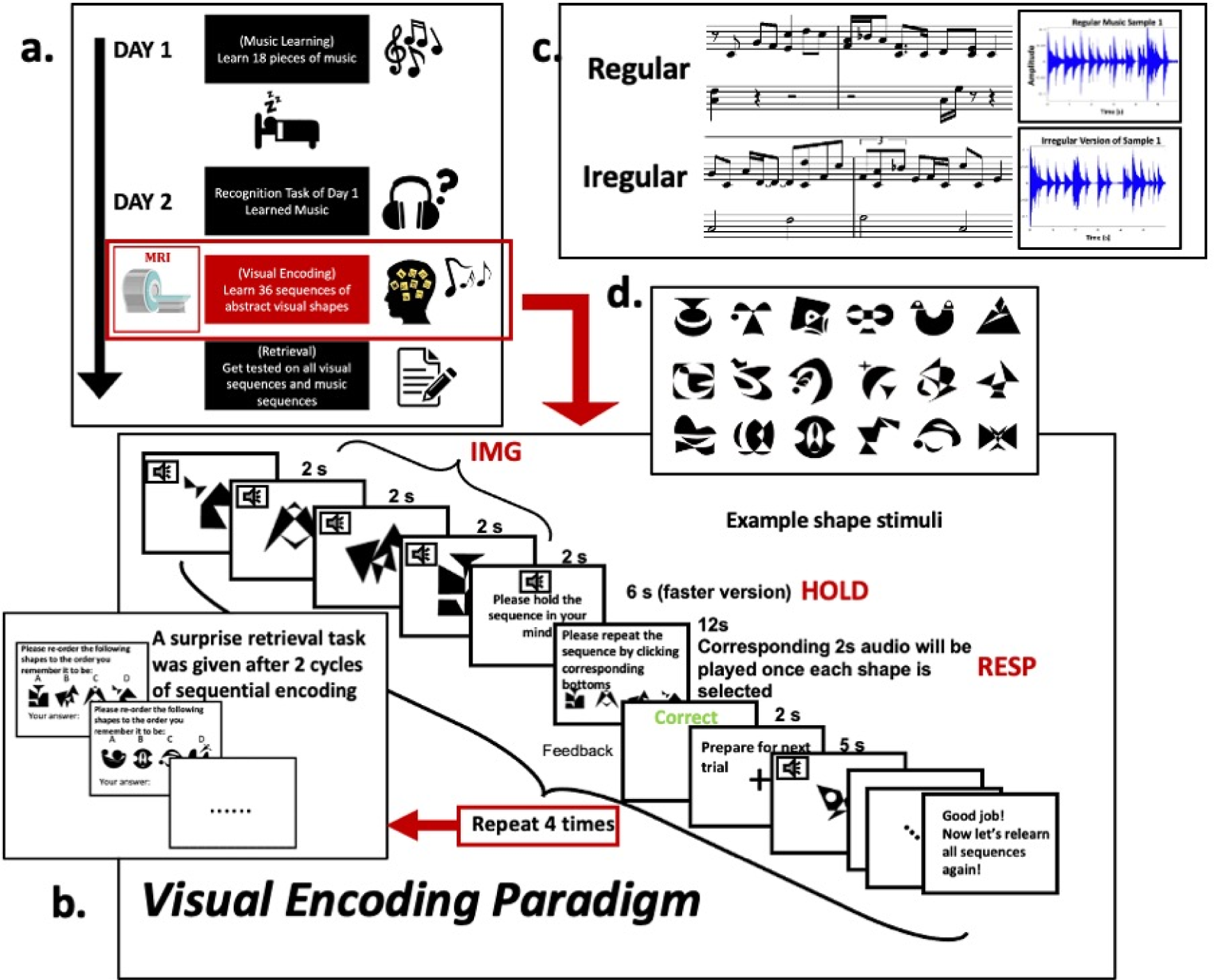
Experimental Design. **a)** two-day experiment: the MRI scan happened during the main visual encoding task **b)** The visual sequences encoding paradigm which happened in an MRI scanner. For each trial, the participants first viewed an image stream and tried to remember the sequence (IMG), then they held the sequence for 6 seconds (HOLD). A retrieval practice question showed up subsequently and asked the participants to retrieve the correct order of the shapes that they just learned (RESP). In the end of each trial, a feedback whether their retrieval was correct or not was given. The encoding contains 4 runs in total. After 2 runs, a surprise retrieval task was given. In the surprise test, the participants needed to retrieve the 36 sequences one by one, without any music or feedback. **c)** example notes and waveforms for regular and irregular music. **d)** example visual stimuli.

To investigate how <u>regularity</u> (schematic/syntactical predictability) of music structure affects concurrent visual sequences learning, we developed two types of music with different levels of syntactical correctness. Specifically, we created 12 pieces of music with syntactically regular structures. Subsequently, we created irregular version of each piece by shuffling the notes and randomizing the temporal intervals between notes (see Figure 1.c for example notes for regular and irregular music). In addition, as a reference, we created isochronous streams serving as the control condition. These monotonic sequences lacked variation across time and were considered the simplest form of music, with the assumption that they would thus not provide temporal cue to parallel visual learning.

For visual stimuli, we created 36 sets/sequences of four shapes (144 shapes in total). To minimize potential semantic associations with real-life objects during visual memory encoding, we made all the shapes novel and abstract, composed of simple lines or curves (see Figure 1.d for example).

To examine the learning effect of the <u>familiarity</u> (veridical predictability) dimension of music predictability, we incorporated a music learning period on the first day of the experiment. On day 1 (training day), participants were instructed to learn a set of 18 music stimuli, randomly selected from the stimuli pool (6 regular, 6 irregular and 6 monotonic). The music learning process spanned approximately two hours and consisted of two main parts. In the first part of the training, participants engaged in repeated listening – they were asked to listen to each piece of music in loops until they felt confident that they recall all the intricate details of the music. The second part of the training involved a music re-composition memory task. For each music piece, we played the first 2 seconds of the music and then provided participants with fragments of the remaining portion, along with some luring choices. Their task was to select and re-order the audio fragments, reconstructing the original music they had learned during the listening phase. This task was performed in loops until participants accurately re-composed all the music without any errors.

On the second day, participants underwent several tasks. The day started with a music recognition task for the music they have learned on previous day. Each trial presented particiapnts with the original music clip and two lure options. Participants were required to select the correct original music clip from the options. The task aim to confirm that participants still remembered the ‘*familiar’* condition music before proceeding to the main task. The monotonic control clips were not included in this recognition test since they were not expected to be actively remembered due to their lack of variation. After the recognition task, participants were brought to the MRI suite for a two-hour scanning session, during which they repeatedly learn 36 sequences of shapes, paired with music. Further details of this task are provided in the next section.

Following the encoding phase in the MRI scanner, participants returned to the lab space to complete two memory retrieval tasks on computers. The first task, music memory task, used an error-detection paradigm. In each trial, participants listened to a version of music that possibly contained difference(s), or errors, compared to the original clip. Their job was to press a bottom whenever they detected an error during the playback. Once the music memory task was completed, participants’ attention was momentarily shifted away from the visual sequences, and they were presented with the final visual memory task. In this task, participants were required to retrieve all 36 sequences they had learned in the scanner without any cues from the music. For each trial, they would see four shapes appeared in random order in a row, their job was to re-order them into the correct order within ten seconds.

### Visual Sequence Encoding

The main task of the experiment, conducted within the MRI scanner, involved visual sequence encoding while music played in the background. Thirty-five subjects participated in the MRI scanning session, while the remaining 16 subjects completed a behavioral version of the encoding task on a computer. In this task, subjects were required to view and memorize sequences of abstract shapes with music. As illustrated in Figure 1.b, for each trial, subjects first viewed a stream of four abstract shapes, with each shape appearing for 2 seconds. Simultaneously, a piece of music was synchronized and played in the background throughout images’ presentation. After viewing the shapes stream, the subjects were required to hold the sequence in their minds for 6 seconds. This served as a buffer before they did a practice retrieval, making the task not too easy. Next, these shapes appeared again in the screen after 6 seconds but in a shuffled order. Subjects’ jobs were to re-order the shapes into the sequence they have seen by clicking the corresponding bottoms on an MRI-compatible response box we provided using their right hands. They had 12 seconds to retrieve the sequence. Once they selected a specific shape, the corresponding 2 seconds of the paired music would play concurrently. We called this ‘<u>Retrieval Practice’</u> period of each trial for encoding. Each trial ended with visual feedback to their retrieval performance, either ‘correct’ or ‘incorrect’.

The encoding task comprised a total of 36 sequences for participants to learn. The entire encoding process was repeated four times, each representing a learning cycle. After completing two cycles of sequential learning, we inserted a surprise retrieval task where we checked subjects’ learning progress. In this task, participants were asked to re-order shuffled shapes into the order they learned without any cues or music. They again had 12 seconds to retrieve each sequence. Each sequence was only tested once. No feedback was provided during the surprise task.

### Behavioral Data Analysis

All behavioral data analyses and visualization were performed in R. We have two main experimental variables: music regularity and music familiarity. For easier analysis, we used a 3×2 factorial design – 3 levels of music regularity which were *control*/monotonic, *irregular* and *regular* and 2 levels of music familiarity. We defined music familiarity as whether the subjects successfully encoded the music on day 1. In the re-composition task on day 1, if the subjects successfully recomposed the music, that piece would be marked as ‘*learned’*. The rest of unsuccessfully re-composed music were marked as ‘*unlearned’* together with music that had not appeared on day 1. We used such strict criterion because we wanted participants to have strong familiarity to ‘learned’ music. Considering individual differences in music skill, individuals might end up with different levels of memory strength for ‘old’ music if we only asked them to listen to music on day 1. To be noted, we also played some monotonic music on day 1. However, due to the nature of no variance across tonality or rhythm, the control music could not be learned technically. For easier visualization and statistical comparison, we labeled the control music appeared on day 1 as ‘*learned’* and the rest as ‘*unlearned’*. To test our main question that how different music structures affect visual sequential learning performance, we aimed on three behavioral measures: 1) during-encoding surprise retrieval task accuracy – this value might indicate visual sequences in which conditions were learned the fastest; 2) the final retrieval accuracy; 3) the response time to retrieve sequences correctly during the final memory task – literatures suggested that a faster reaction time might correlate with a stronger memory(Gimbel & Brewer, 2011). We ran separate generalized mixed model linear regression analyses using *lme*4 package(Bates et al., 2014) to investigate how the fixed effects of music conditions (*regularity*, *familiarity*) and the interaction between them affected each behavioral result, treating subjects as the random effect or the repeated measures. We used a Kenward-Roger approximation(Halekoh & Højsgaard, 2014) and the parametric bootstrap method for getting denominator degrees of freedom and p values for model regressors. For each model, we also used Tukey’s HSD post-hoc test to do pair-wise comparison between different music conditions and we marked all significant differences on the visualizations.

### fMRI Acquisition and Preprocessing

Brain scans were collected using a 3T Siemens Prisma scanner with a 32-channel head coil at the GSU/GT Center for Advanced Brain Imaging. High-resolution T1-weighted structural images were obtained at the beginning using a magnetization prepared rapid gradient echo sequence (MPRAGE) sequences (TR = 2530 ms; TE = 1.69 ms; FOV = 256 * 256 mm; voxel size = 1 mm isotropic; flip angle = 7°). Functional T2*-weighted data and localizer data were obtained following using echo planar imaging (EPI) sequences (TR = 1200 ms; TE =30 ms; FOV = 1540 * 1540 mm; voxel size = 2.5 mm isotropic; flip angle = 65°). The T2*-weighted images provided whole-brain coverage and were acquired with slices oriented parallel to the long axis of the hippocampus. All participants were provided with foam ear plugs and a noise-cancelling MRI-compatible headset. The purpose was to let them be able to talk to the experimenter between runs in the scanner as well as to better listen to the music stimuli during scanning.

All MRI data were preprocessed in the Analysis of Functional Neuro Images software (AFNI: http://afni.nimh.nih.gov/afni/)(Cox, 1996). The preprocessing steps followed AFNI’s standardized pipeline (afni_proc.py). These steps included 1) despiking (*3dDespike*) to reduce extreme values caused by motion or other factors; 2) slice time shifting (*3dTshift*) to correct slice differences in acquisition time; 3) volume registration and alignment (*3dvolreg*) to the volume with the minimal outlier fraction; 4) scaling and normalization of time series into percent of signal change. We used a motion censoring threshold of 0.3mm to remove outlier TRs. All EPI images were warped to MNI templates space using non-linear transformations supported by AFNI’s @SSwarper function. We smoothed all BOLD images using a 4mm full-width at half maximum Gaussian kernel.

### fMRI analysis

In order to investigate how the functional BOLD responses were modulated by different music conditions during visual sequences encoding period, we ran the whole-brain voxel-wise general linear model analysis (*3dDeconvolve*) for each subject. Except for the six nuisance regressors accounting for head motion (translational and rotational) and three accounting for scanner linear drift, the model included fifteen task-related block regressors for five music conditions (*control (C)*, *regular learned (RL)*, *regular unlearned (RU)*, *irregular learned (IL)*, *irregular unlearned (IU)*) at three behavioral events during each trial (*images presentation (IMG)*, *working memory (HOLD)*, *retrieval practice (RESP),* see Figure 1.b for visual paradigm of these three behavioral events). Due to the interaction effects we found from behavioral results, in MRI analysis, instead of two categorical variables (regularity and familiarity), we divided the music into the five conditions to better understand the differences between them in neural responses. The *learned* control and *unlearned* control were combined due to no statistical differences and no technical difference between them. We were specifically interested in how different music affected visual memory encoding, thus we focused on comparing music conditions during *images presentation/initial encoding*, when paired music was played in parallel. We also put strong emphasis on *retrieval practice,* during when we believed any feedback from the paired music might enhance visual sequential encoding. Long-term memory literature suggested that retrieval practice being effective learning strategy and helped acquisition of knowledge even without feedback(Roediger & Butler, 2011). Here we hypothesized that music and its sequential structure might improve the practice retrieval effect and make learning even faster.

To test our hypotheses, on group-level analysis, we ran two separate linear mixed effect models (*3dLME*)(Chen et al., 2013) to compare the neural responses during *IMG* and *RESP* between conditions. We included the subjects as a random effect, considering potential individual differences in how overall music affected them. To reduce family-wise error and use non-parametric comparisons, we applied a Monte Carlo simulation program (*3dClustSim*) with smoothing estimates derived from first-level model residuals (*3dFWHMx*) to get probability of false positive clusters. Based on the result, we reported group-level effects with cluster size larger than 19.3 using a voxel-level threshold of p < 0.001 to determine a cluster-level threshold of corrected p < 0.005 (using 3dClustSim’s second-nearest neighbor clustering option – faces or edges touch). To interpret how music familiarity and regularity interact to modulate visual learning performance, we specifically extracted and visualized neural responses (betas) for clusters that met the criteria above and fell into our regions of interests.

Because we hypothesized that music influences would emerge in part from collaboration between schemas and new learning, we also tested how regions that support different memory processes might together contribute to performance in this cross-modal parallel learning paradigm. Thus, we performed seed-based generalized psychophysiological interaction (gPPI) analysis(McLaren et al., 2012) to test context dependent functional connectivity. Among the brain regions that showed correlated neural responses to music conditions from the univariate GLM results, we were explicitly interested in three brain regions and how their coupling with the rest of brain areas during visual encoding were modulated by our music conditions. These areas were the hippocampus, the prefrontal cortex and the striatum. To generate seed regions, we created 5mm sphere around the peak coordinate of the hippocampus and striatum from the univariate results (anterior hippocampus [L]: MNI –20, –26, –10; dorsal lateral putamen [R]: MNI 28 2 –10). Finally, because prefrontal cortex is a large area and our univariate results revealed separate (albeit neighboring) clusters in the prefrontal cortex, we opted to use the Brainnetome atlas(Fan et al., 2016) embedded in afni to draw a left prefrontal cortex cluster as seed for gPPI analysis, which encompassed the two main superior frontal and superior medial gyrus clusters revealed by the univariate results (size = 1654 voxels). We first extracted the average time series of the seeds (*3dmaskdump*) and removed the trend from the time series (*3dDetrend*). We then generated the interaction term (PPI term) by multiplying the task regressor and the time-series regressors. Lastly, we used *3dDeconvolve* to test the significance of the interaction regressors beyond the main effects and to reveal what brain regions would show differences in its functional connectivity with the seed areas responding to different music conditions. We ran separate generalized psychophysiological interaction (gPPI) general linear models for *IMG* and *RESP* phases. Thus, each model contained five task condition regressors (*C*, *RL*, *RU*, *IL*, *IU*) and the five corresponding PPI regressors plus the time-series regressor and nuisance terms. At the group level, we utilized the same mixed model linear regression and non-parametric multiple comparison correction from above (voxel-level threshold p <0.001, cluster-level threshold p < 0.005, cluster size larger than 21) to identify clusters that showed context-dependent functional connectivity with each region of interests.

## Results

### Behavioral Results

Throughout the experiment, we had two retrieval tasks to record subjects’ memory performance for the visual shapes sequences, one during the encoding after two cycles of sequences learning (surprise task) and one after four cycles of learning (final task). On the surprise task, the overall accuracy ranged from 6.25% to 100% with a mean of 58.13% across subjects (SD = 21.36%). On the final visual sequence retrieval task, participants ranged from 30.56% to 100% in their overall accuracy, with a mean of 83.39% and a standard deviation of 16.86%. We compared the sequences retrieval accuracy between all music conditions versus the control conditions (Figure 2.a). We ran a pairwise t-test and it showed that sequences in the music conditions had significantly higher retrieval accuracy than control conditions during the surprise task (t_50_ = –3.1356, p = 0.001, one sided). No difference was identified between music and control conditions during the final task.

**Figure 2.**
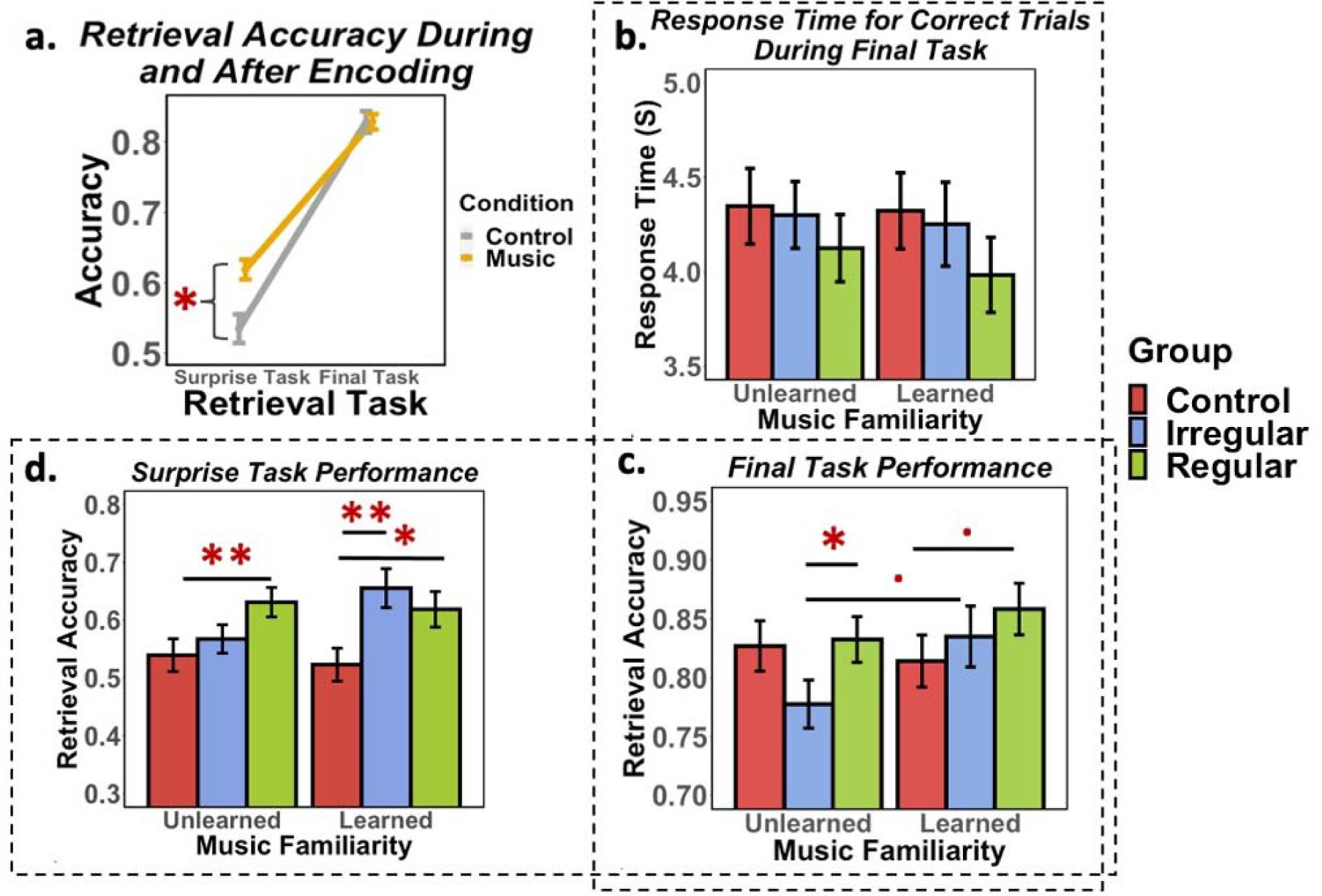
Visual Sequence Memory Performance. (**a**) Visual sequences retrieval accuracy for music conditions versus control condition in the middle of encoding and after encoding. Each dot represents the mean retrieval accuracy across subjects (N=51) for each condition. Error bars represent the SEM. (**b**) Response time for correctly retrieved sequences, comparing learned vs. unlearned conditions and across three regularity conditions. (**c**)(d)Average retrieval accuracy for visual sequences for each group comparing learned vs. unlearned, three regularity conditions. (c) Final retrieval task after encoding. (**d**) surprise retrieval task in the middle of encoding. Tukey’s HSD pair-wise comparisons indicated significant differences between groups: **p<0.001: ***, p<0.01: **, p <0.05: *, 0.05< p<0.1:·**

To further understand how different types of music affected visual sequences encoding, we used a mixed-effect logistic regression model to test how music regularity (control, *irregular*, *regular*) and familiarity (*learned*, *unlearned*) as fixed effects affected whether people could successfully retrieve the corresponding sequence, treating individual subjects as random effect. We first looked at the final retrieval task, for which we found a significant effect of the music regularity (Type 2 ANOVA Wald Chi-Square test: Chi-square = 6.19, df = 2, p = 0.045). Due to a high retrieval accuracy in the final task, we used a pair-wise t-test with unadjusted p value to reveal trending differences between music conditions in the retrieval accuracy. We found *regular* music paired visual sequences were retrieved with more accuracy than *irregular* music in *unlearned* condition (p = 0.04). We also found *learned* music showed a trending effect to increase retrieval accuracy compared to *unlearned* music, in *irregular* condition (p = 0.07).

*Learned regular* music also showed trending better retrieval accuracy than control (p = 0.09). We also looked at response time for correctly retrieved visual sequences during final retrieval task. Memory literature and our prior experiment have used response time or reaction time as a measure of memory strength – a faster retrieval might correlate with more efficient memory recall(Gimbel & Brewer, 2011; Robinson et al., 1997). Using a mixed effect linear regression model, we again identified a significant fixed effect of music regularity (Type 3 ANOVA with Satterthwaite’s method: F =4.00, df_num_ = 2, df_den_ = 1478.2, p = 0.018). Figure 2.b showed that the response time was faster to retrieve a visual sequence when the paired music was regular, in both *learned* and *unlearned* music conditions. Music condition in general showed a faster response time compared to control. We then tested the accuracy between conditions for the surprise task, using the mixed model logistic regression model. It again showed that visual sequences encoded with different music regularity were retrieved with significantly different accuracy during the surprise task (Chi-square = 14.2421, df = 2, p = 0.0008). Pairwise comparisons further suggested that all music conditions, except for *irregular unlearned* music, had higher visual sequences retrieval accuracy than the control (*learned regular*: p = 0.047, *unlearned regular*: p = 0.002; *learned irregular*: p = 0.007).

### fMRI results

#### GLM results for whole brain activity during image presentation and initial encoding

Based on the behavioral results, we mainly focused on two questions when analyzing fMRI data, the first is how music overall modulated brain activity during visual encoding compared to control condition. Considering the behavioral data suggesting that music overall showed an enhancing effect on visual learning, we suspected any neural differences between music and control condition outside music processing related areas might be associated with this behavioral effect. The second question we aimed to investigate in fMRI data was that within music conditions, how different levels of music predictability modulated brain activity and how such differences correlated with any behavioral performance differences.

Under the generalized linear model, we ran t-test contrasts to estimate bold activity differences between all music conditions and control condition (see Table 1). Music conditions showed more activity in the auditory and motor related areas such as inferior frontal gyrus (IFG), superior temporal gyrus (STG), precentral gyrus and cerebellum (Figure 3.a: top). In contrast, music condition revealed decreased activity in parahippocampal gyrus (Figure 3.b.iii). In fact, an exploratory analysis of this effect showed a positive correlation between parahippocampal activity during visual sequences encoding and final retrieval performance in the control condition (r = 0.3, p = 0.044), as one might expect from prior literature tying this area to visual learning, but that correlation was not present for music conditions (r = 0.01, p = 0.89) supporting reduced dependence on its level of engagement (Figure 3.b.i).

**Figure 3.**
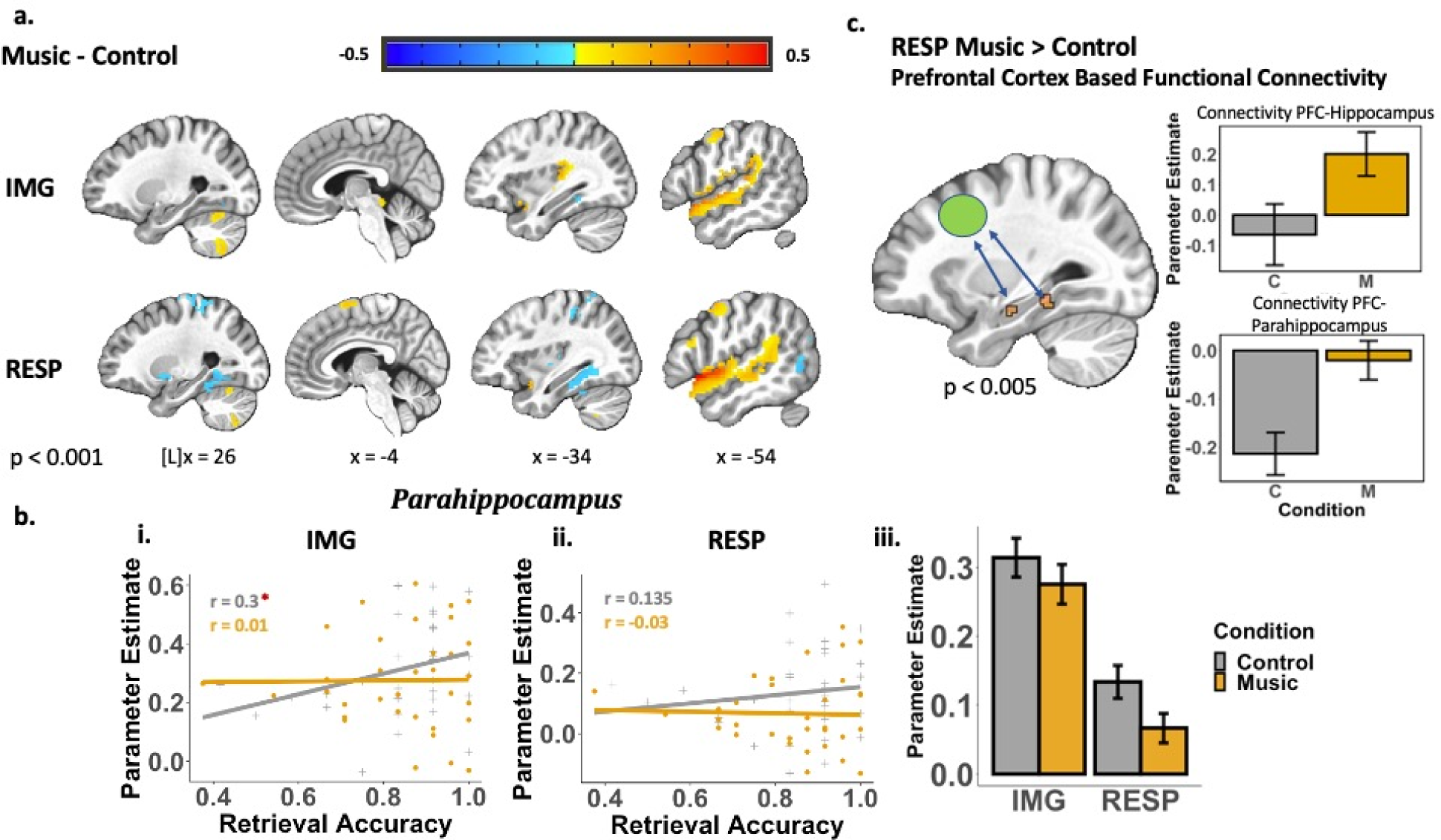
Music and Control Condition Comparisons. [**a**] Whole-brain univariate analysis comparing memory encoding associated brain activity during (1) image presentation (IMG) and (2) retrieval practice (RESP). The sagittal slices displayed significant clusters identified when contrasting music and control (monotonic) conditions. [**b**] (i).(ii). correlation between percent signal change in the parahippocampus and final visual sequences retrieval accuracy during image presentation/initial encoding and retrieval practice, comparing music and control conditions. Each dot stands for one subject. Pearson’s correlation test revealed significant positive correlation between parahippocampal signal with memory performance during initial encoding. (iii). Average percent signal change in parahippocampus with error bars comparing music and control during image presentation/initial encoding(IMG) and retrieval practice(RESP). [c] Generalized psychophysiological interaction(gPPI) analysis using prefrontal cortex as seed regions. Hippocampus and parahippocampus both showed stronger functional connectivity with PFC in music conditions contrasting control during retrieval practice (RESP).

**Table 1.** Whole brain voxel based analysis results during image presentation (IMG)

| Cluster Size (Voxel) | Peak Coordinates(RAI) |  |  | z-score<br>(2-sided) | Regions |
| --- | --- | --- | --- | --- | --- |
|  | x | y | z |  |  |
| During Image Presentation / Initial Encoding |  |  |  |  |  |
| Control > Music |  |  |  |  |  |
| 24 | 2.5 | 39.2 | -9.5 | -3.66 | Parahippocampus |
| 20 | -32.5 | 39.2 | -14.5 | -3.83 |  |
| Music > Control |  |  |  |  |  |
| 1113 | -50 | -5.8 | -7 | 10.52 | Superior Temporal Gyrus |
| 983 | 50 | -0.8 | -4.5 | 11.41 | Inferior Frontal Gyrus |
| 36 | -57.5 | 1.8 | 48 | 5.94 | Precentral Gyrus (R) |
| 134 | 22.5 | 71.8 | -59.5 | 4.28 | Cerebellum (L) |
| Learned > Unlearned |  |  |  |  |  |
| 49 | -5 | 39.2 | 38 | 3.74 | Middle Cingulate Gyrus |
| 33 | 7.5 | 41.8 | 38 | 4.34 |  |
| 42 | -27.5 | -35.8 | 50.5 | 3.56 | Superior Frontal Gyrus (R) |
| 22 | -5 | -38.2 | 45.5 | 3.78 | Superior Medial Gyrus (R) |
| Regular > Irregular |  |  |  |  |  |
| 57 | -32.5 | 24.2 | -24.5 | 3.9 | Fusiform Gyrus +<br>Parahippocampus |
| 28 | 32.5 | 34.2 | -22 | 4.47 |  |
| 52 | -27.5 | 79.2 | 15.5 | 3.68 | Middle Occipital Gyrus (R) |
| 51 | 52.5 | 61.8 | -12 | 3.65 | Inferior Temporal Gyrus (L) |
| Irregular > Regular |  |  |  |  |  |
| 103 | 0 | 21.8 | 43 | 3.98 | Middle Cingulate Gyrus |
| 143 | -2.5 | 40.8 | 15.5 | 3.88 | Anterior Cingulate Gyrus (R) |
| 102 | 35 | 86.8 | -37 | 3.5 | Cerebellum (L) |
| 69 | -15 | -38.2 | 50.5 | 3.5 | Superior Frontal Gyrus (R) |
| 65 | -50 | -23.2 | 40.5 | 4.1 | Middle Frontal Gyrus(R) |

Within music conditions, when contrasting *regular* and *irregular* music, results indicated increased activity in inferior temporal gyrus and visual cortex, including fusiform gyrus and middle occipital gyrus, while there was decreased activity in cerebellum, superior/middle frontal gyrus and middle cingulate gyrus in *regular* music compared to *irregular* (igure 4.a). For the familiarity contrast, *learned* music showed stronger activity in superior frontal gyrus, superior medial gyrus and middle cingulate gyrus compared to *unlearned* music.

#### GLM results for whole brain activity during retrieval practice and re-encoding

Comparing all music conditions with control for bold activity during *retrieval practice* of encoding period revealed several significant regions. Once again, we observed increased activity in music related areas for music conditions, including IFG, STG, motor cortex, cerebellum and supramarginal gyrus (igure 3.a: bottom). Meanwhile, music conditions were associated with decreased activity in parahippocampus (Figure 3.b.iii) as well as in the putamen, inferior temporal gyrus, postcentral gyrus, precuneus and visual cortex (Figure 3.a: bottom, see Table 2).

**Table 2.** Whole brain voxel based analysis results during retrieval practice (RESP)

| Cluster Size (Voxel) | Peak Coordinates(RAI) |  |  | z-score<br>(2-sided) | Regions |
| --- | --- | --- | --- | --- | --- |
|  | x | y | z |  |  |
| During Retrieval Practice / Re-encoding |  |  |  |  |  |
| *Control > Music* |  |  |  |  |  |
| 201 | -32.5 | 46.8 | -7 | -4.42 | Parahippocampus |
| 134 | 27.5 | 51.8 | -7 | -4.15 |  |
| 45 | 30 | 14.2 | -2 | -4.02 | Putamen (L) |
| 59 | 37.5 | 39.2 | 58 | -3.45 | Postcentral Gyrus |
| 32 | -37.5 | 34.2 | 48 | -4.41 |  |
| 182 | 47.5 | 51.8 | -22 | -3.65 | Inferior Temporal Gyrus (L) |
| 69 | -45 | 79.2 | 15.5 | -3.63 | Middle Occipital Gyrus |
| 37 | 30 | 89.2 | 30.5 | -3.51 |  |
| 33 | -22.5 | 64.2 | 25.5 | -3.57 | Precuneus (R) |
| *Music > Control* |  |  |  |  |  |
| 1452 | -52.5 | -18.2 | -7 | 9.09 | Superior Temporal Gyrus |
| 635 | 52.5 | -5.8 | -4.5 | 11.38 | Inferior Frontal Gyrus |
| 60 | -57.5 | 1.8 | 48 | 6.22 | Precentral Gyrus |
| 33 | 55 | 4.2 | 53 | 6.49 |  |
| 116 | 62.5 | 44.2 | 25.5 | 5.92 | Supramarginal Gyrus (L) |
| 60 | 30 | 66.8 | -52 | 5.63 | Cerebellum (L) |
| 34 | -5 | -5.8 | 70.5 | 4.7 | Supplementary Motor Area (R) |
| *Learned > Unlearned* |  |  |  |  |  |
| 34 | -27.5 | 14.2 | -22 | 4.15 | Hippocampus (R) |
| 457 | -60 | 46.8 | 48 | 3.76 | Inferior Parietal Lobule |
| 34 | 57.5 | 49.2 | 53 | 4.33 |  |
| 277 | -45 | -23.2 | 40.5 | 4.31 | Middle Frontal Gyrus |
| 27 | 30 | -38.2 | 28 | 3.56 |  |
| 80 | 25 | 89.2 | -24.5 | 3.85 | Cerebellum |
| 56 | -52.5 | 66.8 | -37 | 4.74 |  |
| 222 | -40 | 14.2 | -4.5 | 3.96 | Insula + Striatum |
| 23 | 37.5 | -8.2 | -14.5 | 3.55 |  |
| 213 | -2.5 | 1.8 | 38 | 3.39 | Middle Cingulate Gyrus (R) |
| 57 | -55 | -25.8 | -7 | 3.87 | Superior Temporal Gyrus |
| 31 | 42.5 | 11.8 | -9.5 | 3.88 | Inferior Frontal Gyrus |
| *Regular > Irregular (only within learned conditions)* |  |  |  |  |  |
| 31 | 37.5 | 6.8 | 63 | 4.14 | Precentral Gyrus (L) |
| 26 | 47.5 | 49.2 | -14.5 | 3.47 | Inferior Temporal Gyrus (L) |
| *Irregular > Regular (only within learned conditions)* |  |  |  |  |  |
| 19 | 32.5 | 41.8 | 0.5 | -4.38 | Hippocampus (L) |
| 63 | -5 | 89.2 | 28 | -4.31 | Cuneus |
| 50 | 2.5 | 89.2 | 23 | -4.32 |  |
| 64 | -20 | 41 | 64 | -3.46 | Postcentral Gyrus (R) |

No clusters survived cluster threshold correction when comparing regular and irregular conditions during *retrieval practice*. When examining the subconditions during retrieval practice, comparing regular and irregular music within learned and unlearned conditions, we found no differences between regular and irregular conditions under unlearned conditions. However, when focusing on learned music conditions alone, such that familiarity effects were less likely to dominate signal differences, we demonstrate that inferior temporal gyrus and precentral gyrus were more active during retrieval practice when the task was paired with regular (learned) music than irregular. Conversely, we found hippocampus, cuneus and postcentral gyrus were more active in irregular (learned) music conditions (Table 2).

When contrasting *learned* versus *unlearned* conditions, we identified more BOLD activity in IFG, STG, cerebellum, inferior parietal lobule as well as in the hippocampus, middle frontal gyrus, middle cingulate cortex and insula (Table 2). Further pair-wise comparison on hippocampal BOLD activity revealed decreased hippocampal activity in learned music compared to both control and unlearned during both *image presentation* and *retrieval practice* (Figure 5.a).

#### Functional Connectivity (gPPI)

As described in the methods, we targeted three regions from the univariate fMRI analyses to run generalized psychophysiological interaction analysis in order to investigate interaction and collaboration between brain regions, based on their respective functional relevance in the prior literature for music-guided memory (as detailed in the introduction) and their interconnected nature potentially relevant for sequencing across a broad class of memory paradigms (e.g. Brown et al., 2012).

First, for our striatal (putamen) seed, <u>during image presentation (*IMG*)</u>, there were no connectivity differences between music and control. However, when comparing *regular* and *irregular* conditions, results revealed putamen had stronger functional connectivity in the *irregular* condition with cerebellum (# of voxel = 52, [-25, 39, 49.5]), middle cingulate gyrus (# of voxel = 31, [-2.5, 16.8 35.5]) and right middle frontal gyrus (# of voxel =25, [-32.5, –15.8, 58]), whereas its connectivity was stronger in the *regular* condition with the left anterior hippocampus (# of voxel = 23, [25, 14.2, –12]) (Figure 4.b). T-tests investigating the effect of music familiarity revealed putamen had stronger connectivity with a broad set of broad cortical areas in *learned* condition, including both hemispheres’ supplementary motor area (SMA) (# of voxek = 491, [1,19.2, 50.5]), right middle frontal gyrus (# of voxel = 169, [-45, –3.2, 58]), left precentral gyrus (# of voxel = 57, [32.5, 6.8, 60.5]) and right superior parietal lobule (# of voxel = 49, [-42.5, 51.8, 58]). <u>During retrieval practice (*RESP*)</u>, t-tests demonstrated stronger connectivity between putamen and visual cortex (# of voxel = 55, [12.5, 79.2, 10.5]; cluster size = 36, [-12.5,76.8, 13]) in *regular* compared to *irregular* condition. Putamen also had stronger connectivity with entorhinal cortex (# of voxel = 48, [-35, 14.2, –27]) in the *unlearned* condition compared to the *learned* during retrieval practice (Figure 5.b).

**Figure 4.**
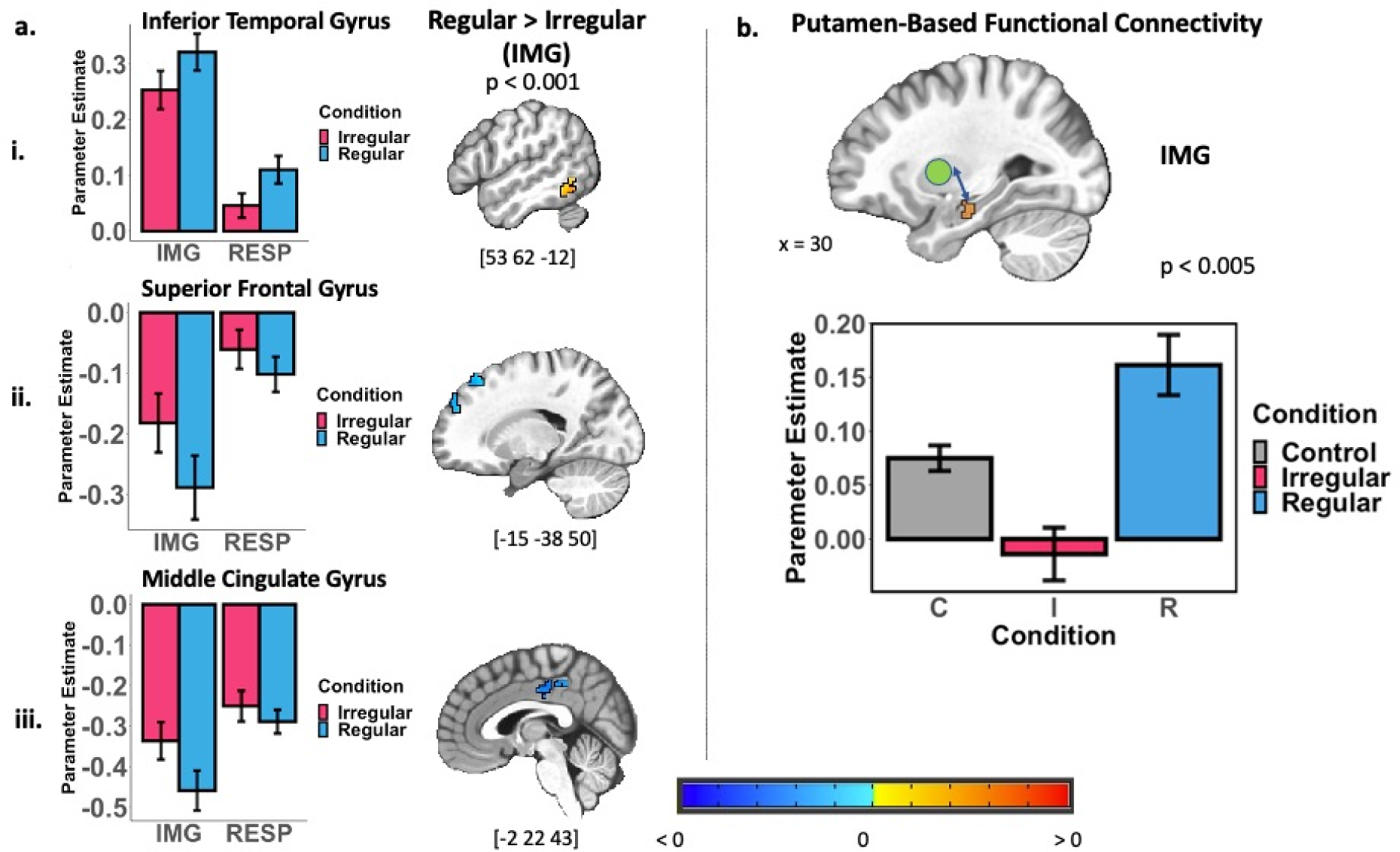
Regular and Irregular Music Conditions Comparisons. [**a**] Whole-brain univariate analysis of memory encoding associated brain activity comparing regular and irregular music conditions. The sagittal slices displayed significant clusters identified during image presentation/initial encoding (IMG). Regions with different activation during visual encoding with regular versus irregular music included but not limited to inferior temporal gyrus, superior frontal gyrus and middle cingulate gyrus. Bar plots for each region displayed average percent signal change in the two music conditions. [**b**] Generalized psychophysiological interaction (gPPI) analysis using putamen as seed regions during retrieval practice. Right anterior hippocampus showed stronger functional connectivity with putamen in regular music conditions contrasting irregular music during image presentation and initial encoding (IMG). Bottom: the bar plot showed average parameter estimate of the putamen-anterior hippocampus functional connectivity for different music conditions during image presentation.

**Figure 5.**
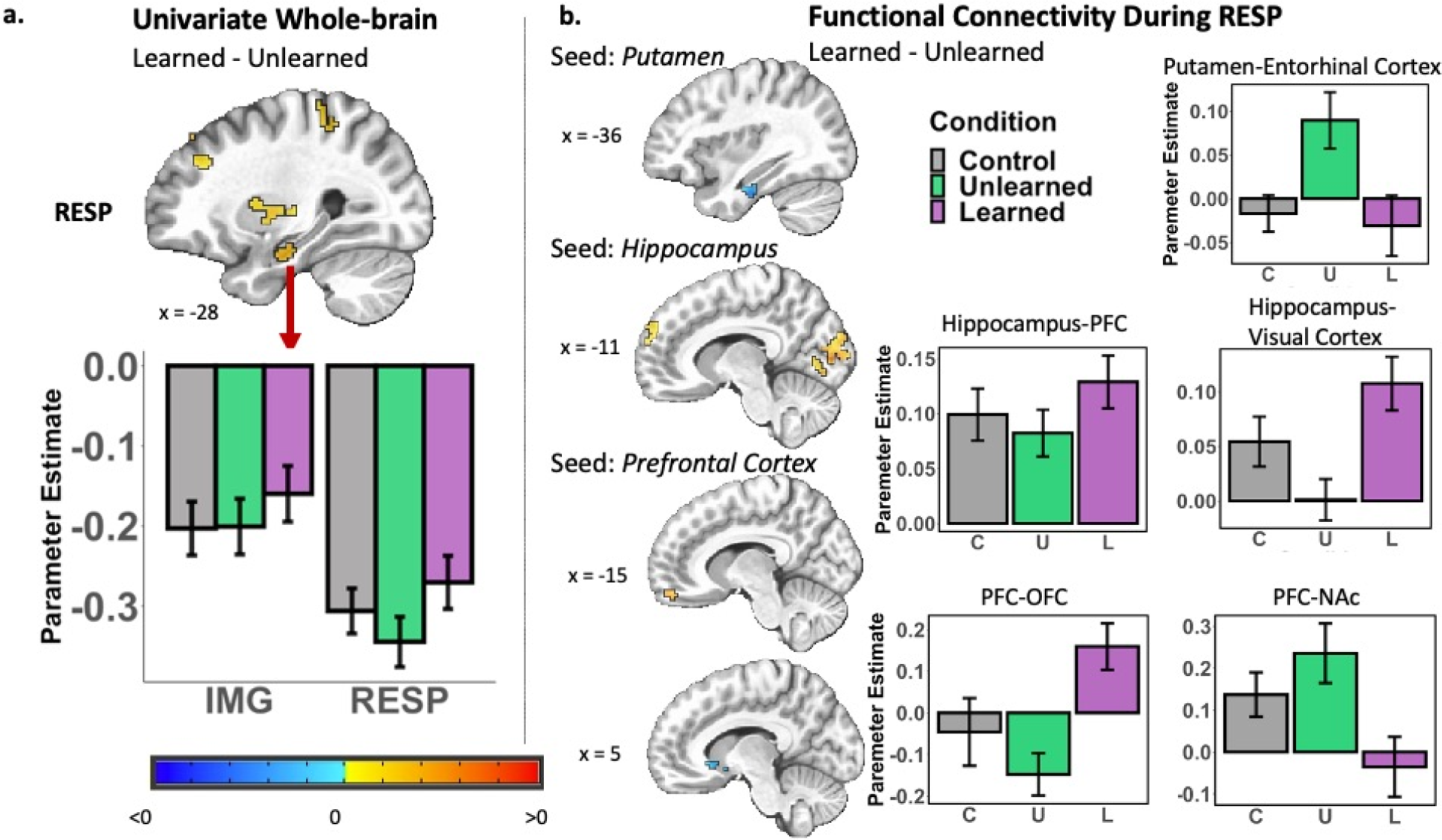
Learned and Unlearned Music Conditions Comparisons. [**a**] Whole-brain univariate analysis of memory encoding associated brain activity comparing learned and unlearned music conditions. The sagittal slices displayed significant clusters identified during retrieval practice (RESP). Regions with different activation during retrieval practice with learned versus unlearned music included but not limited to the hippocampus, middle frontal gyrus and SMA. Bar plot displayed average percent signal change in the two music conditions in the hippocampus, one of our regions of interest. [**b**] Generalized psychophysiological interaction (gPPI) analysis using our three regions of interest (Putamen, hippocampus and prefrontal cortex) as seed regions during retrieval practice. Most significant networks were found during retrieval practice (RESP). The sagittal slices displayed main networks identified to be differently involved in retrieval practice for learned vs. unlearned conditions. These main networks we were interested at included: putamen-entorhinal cortex, hippocampus-visual cortex, hippocampus-prefrontal cortex (peak at the superior frontal gyrus), prefrontal-orbital prefrontal cortex and prefrontal-nucleus accumbens. Bar plots displayed the average parameter estimate of the PPI regressor of each music conditions during retrieval practice.

The second region we ran seed-based gPPI from was the hippocampus. <u>During image</u> <u>presentation (*IMG*)</u>, when contrasting music versus control or regular versus irregular conditions, no differences in functional connectivity were identified with any regions and right hippocampus. However, when examining the effect of music familiarity, visual cortex (# of voxel = 80, [0, 84.2, 5,5]) and right supramarginal gyrus (# of voxel = 21, [-55, 44.2, 43]) demonstrated stronger connectivity with right hippocampus in the *learned* condition. By contrast, hippocampal and caudate (# of voxel = 40, [7.5, 18.2, 18]) connectivity was stronger in *unlearned* conditions. <u>During retrieval practice (*RESP*)</u> when examining music regularity, right hippocampus had reduced connectivity with left middle temporal gyrus (# of voxel =34, [45, 61.8, –2]), left SMA ((# of voxel =28, [0, 11.8, 48]) and left precuneus (# of voxel = 21, [10, 54.2, 43]) in the *regular* condition. When comparing *learned* and *unlearned*, cerebellum (# of voxel = 88, [12.5, 54.2, – 34.5]), visual cortex ((# of voxel = 78, [-10, 84.2, 3]) and left superior frontal gyrus (# of voxel = 53, [22.5, –63.2, 13]) exhibited stronger hippocampus coupling in the *learned* condition than the *unlearned*.

Finally, the third seed region of interest was the prefrontal activity volume shown to be modulated by music familiarity and regularity in the univariate results. <u>During image</u> <u>presentation period (IMG)</u>, there were no brain regions with differential connectivity with this seed between any contrasts. However, <u>during retrieval practice (RESP)</u>, the *regular* versus *irregular* contrast was characterized by reduced connectivity for *regular* between the seed and right superior temporal gyrus (# of voxel = = 125, [-47.5, 44.2, 18]), right middle temporal gyrus (# of voxel =42, [-57.5, 9.2, 18]), right middle cingulate cortex (# of voxel = 37, [-2.5, 26.8, 48]) and the parahippocampal gyrus (# of voxel size = 21, [20, 41.8, –14.5]). When contrasting the *learned* and *unlearned* conditions, *learned* exhibited stronger connectivity between PFC and right orbital prefrontal cortex (# of voxel = 36, [-15, –55.8, –19.5]) as well as between PFC and left nucleus accumbens (# of voxel = 26, [5, –5.8, –9.5]).

To summarize, one important theme of the gPPI analyses synthesized in Figure 5 is that retrieval practice was characterized by elevated striatal connectivity for *unlearned* (or, even decreased striatal connectivity for *learned* relative to control) with prefrontal and MTL regions (putamen, nucleus accumbens [NAc] effects), whereas prefrontal-hippocampal interconnectivity showed the opposite pattern. This was of particular interest because retrieval practice was the time window in which neural processes relating the tones heard during sequence retrieval to prior knowledge of the music can provide an explicit feedback pathway for the visual sequence memory.

## Discussion

This study employed fMRI to investigate how music of varying predictability levels influenced concurrent visual sequential encoding. We found that listening to predictable music reduced engagement in areas associated with long-term sequential memory (like the MTL and striatum), as well as areas associated with working memory and attention in the PFC. We also observed altered connectivity among these brain areas during encoding and retrieval practice depending on the music conditions. We theorize that the music enhancing effects are brought about by a combination of mechanisms, such as attention modulation, schema processing and reward effects, all intricately tied to the functions of these subdivisions and their interactions.

Our data showed that visual sequences encoded with music of greater predictability can be learned better (indicated by the surprise test and final memory task). Comparing music with monotonic control, we observed decreased engagement of parahippocampal cortex, a critical region associated with visual associative memory (Hayes et al., 2007). We also found decreased activity in areas extending to putamen and SMA, which have been strongly associated with temporal(Macar et al., 2002), sequential(Rauch et al., 1997), and grammar (Tagarelli et al., 2019) learning. These results suggest that when music is present, sequential learning may become more efficient, thereby reducing processing demands on temporal sequencing systems.

Previous studies have indicated that retrieval practice immediately after encoding facilitates recollection (Roediger & Butler, 2011), and is supported by working memory areas including PFC and SMA(Barbey et al., 2013; Cañas et al., 2018; Curtis & D’Esposito, 2003). Additionally, numerous studies have indicated parahippocampal gyrus’s critical role in visual memory formation(Eichenbaum & Lipton, 2008), particularly in binding information (Hsieh et al., 2014) during initial encoding and working memory(Luck et al., 2010). Alongside our evidence for increased frontal engagement and reduced MTL and striatal engagement during encoding and retrieval-practice with music, we suggest the decreased correlation between parahippocampal activity and memory success offers additional evidence that music’s predictable sequential structure either enhanced established sequence encoding mechanisms or perhaps supported a to-be-determined alternative form of encoding necessary sequence information outside the visual ventral stream, sharing the processing burden(Kravitz et al., 2013).

One possible mechanism for more efficient learning with paired music could be facilitated working memory, potentially through attention modulation. Our behavioral data revealed that regular music induced significantly greater enhancement than irregular music on parallel encoding; prior literature has emphasized the importance of music regularity, especially the rhythmic predictability, for music enhancement of cognitive functions (Nguyen & Grahn, 2017; Thompson et al., 2001). Studies suggest the temporal expectation of background music could direct attention to specific timing features (Basu & Banerjee, 2022; Bolger et al., 2013), and in our design *irregular* music would disrupt this by rendering the music syntactically-unpredictable. Notably, during encoding in *regular* music conditions, we observed decreased activity in default mode network (DMN) related areas such as dorsolateral and dorsomedial prefrontal cortex as well as middle cingulate cortex (Table 1; Figure 4) – deactivation in DMN correlates with working memory and attention (Mayer et al., 2010). Furthermore, we found decreased activity in cuneus, a region involved in modulating visual spatial memory and attention (Simpson et al., 2011), in *regular* music during retrieval practice, also consistent with any reduced attentional demands or workload needed during retrieval practice for this condition. Collectively, these findings support the notion of improved efficiency in visual attention and working memory streams for visual sequences encoding during *regular* music listening, while complemented by *stronger* activation in visual information processing areas such as inferior temporal gyrus (Figure 4) in *regular* encoding and retrieval practice. We also observed stronger connectivity between the putamen and anterior hippocampus during encoding for *regular* conditions. Putamen is particularly important for temporal rule learning (Nenadic et al., 2003) while anterior hippocampus has long been established as crucial for encoding new associative memories(Chua et al., 2007). Heightened communication between these two areas could contribute to the enhanced efficiency encoding temporal and relationships of the visual stimuli, observed in the behavioral data.

However, we suspect there are other putative mechanisms besides improved attention behind music enhancements. One reason is that our control isochronous condition also provides low-level predictability (regular rhythm) that some studies have suggested can improve attention(Escoffier et al., 2010). Yet, in our design most music conditions, even the *learned irregular* condition, showed better visual learning results than control. This discrepancy suggests that the benefits from music may relate to mechanisms beyond attention enhancement.

One possibility is schema processing: in which prior knowledge influences how new information is learned. Music is highly schematic and can provide a rhythmic ‘scaffold’ for new sequence learning(Conway et al., 2009; Leman, 2012). Studies have shown stronger connectivity between PFC and MTL during encoding if to-be-learned information is associated with prior schemas, with the PFC aiding in accessing old memories, projecting new information onto the “schema” and reducing encoding demands on the MTL (van Kesteren et al., 2012). Although our behavior did not reveal a significant main effect of familiarity, we observed significant differences between *learned* and *unlearned* conditions for *irregular* music. The improvement in *learned irregular* music indicates that music familiarity can lead to memory benefits and can overcome the irregularity of the music. We also identified neural differences between the two levels of familiarity during visual sequential learning, potentially explaining the faster response time in *learned* conditions (Figure 2.d) and the beneficial effect from *learned irregular* music (Figure 2.c). Specifically, during initial encoding, we found increased activity in superior frontal areas and cingulate gyrus (Table 2). These results align with music literature suggesting that familiar music recognition is supported by superior frontal gyrus(Freitas et al., 2018) and could induce familiarity-based emotional reactions via cingulate gyrus function(Pereira et al., 2011). During retrieval practice, we observed distinctive *learned* versus *unlearned* differences outside music recognition areas. Of note, hippocampus was significantly more active with *learned* music. The hippocampus is not frequently reported in music memory tasks, where extant data have argued it mainly supports declarative memory encoding(Esfahani-Bayerl et al., 2019; Tulving & Markowitsch, 1998). Stronger hippocampal activity in our data may support enhanced encoding of visual information, mediated by processing elsewhere of the music and its structure. In fact, our data revealed stronger connectivity between PFC and MTL (both hippocampus and parahippocampal cortex) specifically during retrieval practice, when the participants received music as a form of feedback when re-ordering the shapes (corresponding audio played for each selected shape). During this time window, the sequential structure of familiar music, putatively accessed by PFC(Freitas et al., 2018), may serve two purposes: a) detecting errors and b) guiding continued re-encoding of visual sequences via MTL. Moreover, we found stronger connectivity for *learned* music conditions between putamen, a subcortical temporal structure processing area(Nenadic et al., 2003), and the entorhinal cortex, the gateway into the hippocampus (Eichenbaum & Lipton, 2008). Some studies have proposed that such striatal-MTL connectivity facilitates processing temporal expectations for associative memory(van de Ven et al., 2020). In our case, this increased connectivity is consistent with the fact that familiar music provides the most robust temporal expectation or “template” for visual sequential encoding.

A third (not mutually exclusive) framework for the benefits of familiar music in our design is reward processing. Music that follows predictions has been shown to activate reward circuitry including the ventral striatum (Pereira et al., 2011; Salimpoor et al., 2013). Reward signaling can promote learning via dopaminergic plasticity modulation(Wittmann et al., 2005). Prefrontal cortex, ventral striatum and hippocampus may jointly regulate reward-based learning in humans(O’doherty, 2004; Schott et al., 2008), and our data revealed interactions between these areas for *learned* music: we found elevated hippocampal and ventral caudate connectivity during initial encoding for *learned* music conditions, while prefrontal cortex, by contrast, showed reduced connectivity with ventral striatum during retrieval practice for *learned* conditions. While we can only speculate how these differing network shifts modulate familiar music effect during visual sequential learning, our data align with theoretical frameworks which use reward signaling from music to account for learning(Gold et al., 2019). We aim to explore how music predictability guides reward feedback during concurrent learning in our future work.

In summary, our study demonstrates a positive impact of predicative music on concurrent visual sequential/associative learning. The data suggest more-efficient encoding during predictable music listening, evidenced by decreased activity in sequence learning areas including parahippocampal cortex and striatum. Regularity effects were particularly prominent during initial encoding, possibly linked to improved working memory and attention performance and decreased PFC activity. Familiarity, on the other hand, could provide a known temporal structure that aids novel sequence learning, and this significantly altered hippocampal-frontal-strialtal interactions especially during retrieval practice. Given the widespread habit of (and interest in) music listening during various cognitive activities, it is our hope that understanding what types of music offer cognitive benefits might inspire real life music design and uses in educational, work, and clinical settings.

## Acknowledgments

This work was supported in part by the National Institute on Aging of the National Institutes of Health under Award 1-R21AG063131.

